# Mitochondrial hitch-hiking of *Mapt* mRNA maintains Tau levels in axons

**DOI:** 10.64898/2026.01.09.698691

**Authors:** Hariharan Murali Mahadevan, Ayat E. Rehailia, Adrian Marti Pastor, Isabel M. C. Geelhaar, Luciano Román-Albasini, Regina Feederle, Monika S. Brill, Thomas Misgeld, Angelika B. Harbauer

## Abstract

Hitch-hiking of transcripts on organelles, including mitochondria, has emerged as a common pathway to transport mRNAs into the axon to enable their local translation. However, the extent of mitochondrial mRNA association *in vivo* has not been established. Here we report the landscape of mitochondrial mRNA association in two nerve tissues, the retinal ganglion cell derived optic nerve and the motor neuron derived part of the sciatic nerve. This revealed astonishing tissue diversity between mitochondria associated mRNAs that can partly be explained by the lack of expression of the mitochondrially anchored RNA-binding protein SYNJ2a in motor neurons. This diversity led us to discover that also *MAPT* mRNA, encoding the axon-enriched microtubule associated protein Tau, is transported via the mitochondrial RNA anchoring complex SYNJ2a/SYNJ2BP into axons of glutamatergic neurons. Loss of SYNJ2BP prevents axonal localization of *MAPT* and miss-sorting of Tau, consequently stunting axonal growth in cultured hippocampal neurons. This connects mitochondrial transport to the maintenance of the local cytoskeleton and axonal growth in glutamatergic neurons.

## Main text

Local translation of proteins shapes the proteome of neuronal subcompartments in order to maintain their polarization^1,2^. Among the most abundant transcripts enriched in axonal transcriptomes are those encoding mitochondrial and ribosomal proteins^3–5^, but also evidence for local translation of cytoskeletal elements has been provided. This includes the local translation of *beta-actin* mRNA in axonal growth cones^6^ and at axonal branch sites^7–9^ and the local translation of the axon-specific microtubule-associated protein Tau (MAPT)^10,11^. Tau is an axon-specific microtubule binding protein and thus must be sorted into the axon and removed from the somatodendritic compartment. Pathological missorting occurs during neurodegeneration and is thought to be an early marker preceding the formation of Tau tangles^12^. Both mRNA and protein-based mechanisms have been proposed, and most likely a concert of all these are necessary to support the exclusive axonal localization of Tau in healthy neurons^13^. An HuD-binding site in its 3’UTR has been suggested to mediate its axonal localization^10^, but also binding to the neuronal RNA-binding protein (RBP) heterogeneous nuclear ribonucleoprotein R (hnRNP R) has been observed to contribute to axonal Tau localization^14^. Local translation of Tau during axon development is driven by the increased activity of mTOR signaling due to a TOP motif in its 5’UTR^15^. Tau dependent stabilization of microtubules is suggested to favor axon growth^16^ and to regulate axonal transport^17^. Various isoforms of Tau are expressed during neuronal development and also differ between the central and peripheral nervous system^18,19^.

Interestingly, hotspots of local translation in axons have been reported to form near mitochondria^7,20^ and mitochondria may even serve as means of transport for mRNAs^21,22^. We have recently shown that the complex composed of SYNJ2BP (Synaptojanin 2 binding protein, an outer mitochondrial membrane protein) and SYNJ2a (a splice variant of the RNA-binding protein Synaptojanin 2) anchors the transcript encoding PINK1 (PTEN-induced kinase 1, a mitochondrial quality control protein) to mitochondria, both in hippocampal neurons *in vitro* and in retinal ganglion cells *in vivo* ^21^. Parallel work by the Perlson and Arava labs showed a similar mitochondrial hitch-hiking for the *Cox7c* mRNA in cultured motor neurons^22^. We therefore set out to characterize the full extent of mitochondrial mRNA association in retinal ganglion cells and motor neurons *in vivo*.

### Landscape of mitochondrial RNA association in axons

To isolate mitochondria specifically from the axonal comparment *in vivo*, we took advantage of the recently published MitoTag mouse line^23^, which allows the Cre-dependent expression of a GFP-tag targeted to the mitochondria outer membrane (OMM-GFP). We crossed these mice to ChAT-Cre mice to label cholinergic lower motor neurons and to RBP4-Cre mice to label the glutamatergic retinal ganglion cells (Fig. 1a). We then isolated GFP-positive mitochondria by immunocapture from the optic nerves (ON) and sciatic nerves (SN) of these mice, as these tissues contain the axons of the OMM-GFP expressing neurons (Fig. 1B, labelled GFP). As a comparison, we also isolated all mitochondria from both tissues using immunocapture of Tom22, a universally expressed mitochondrial outer membrane protein (Fig. 1b, labelled Tom). We analyzed the polyadenylated transcripts co-isolated with both mitochondrial isolations by next generation sequencing, as well as the bulk RNA content of the respective tissue (Fig. 1b). After extracting the differentially expressed transcripts for all three possible comparisons (GFP vs. TOM vs. Bulk), we observed a high correlation between transcripts isolated via GFP and TOM isolations normalized to their abundance in the bulk transcriptome (R^2^= 0.609 (ON) and 0.62 (SN), respectively, Fig. S1a-b). Since mitochondria contain their own DNA that produces polyadenylated transcripts in mice, these transcripts served as our positive control as they will naturally be enriched upon isolation of intact mitochondria. Indeed, we observed in both optic and sciatic nerve a significant enrichment for most mitochondrially encoded mRNAs (Fig. S1a-b). The enrichment of mitochondrially encoded mRNAs slightly skewed towards Tom22-isolated mitochondria, in line with recent reports that some axonal mitochondria lack mitochondrial DNA^24,25^. Focussing on the axon-specific isolations, we detected 1853 (ON) and 974 (SN) transcripts associated with mitochondria in both tissues, with 231 shared between both cell types (Fig. 1c). Gene ontology enrichment mapping revealed that the shared mitochondria-associated transcripts formed four functional units: (i) oxidative phosphorylation, driven mainly by the mtDNA-encoded transcripts, (ii) protein modification, (iii) RNA processing and (iv) memory (Fig. 1d). The third category was rather unexpected, as RNA splicing is mainly a nuclear process. However, there is some evidence for tRNA processing near mitochondria^26,27^, fitting to their role as hot-spots for local translation. Interestingly, the relatively small overlap suggested that the association of transcripts to mitochondria may be differentially regulated between tissues. This was particularily evident in the principal component analysis of the RNAseq (Fig. S1b). The first component captures the variability coming from the isolation method, whereas the second component is mostly explained by the tissue from which the isolation was performed. We therefore decided to continue our analysis separately for the two tissues and plotted the transcripts enriched on axonal mitochondria separately for ON and SN isolations (Fig. 1e-f and S1c-f). Again, mtDNA-encoded RNAs were among the most enriched transcripts in GFP and Tom isolations compared to bulk. Strikingly, when we plotted for nuclear-encoded mitochondrial transcipts, only a select few transcripts were significantly associated with mitochondria in axons, 8 in the ON and 3 in the SN. This suggests that the majority of transcripts associating with mitochondria strongly enough to survive mitochondrial isolation are not providing proteins destined to work in mitochondria. Additionally, of the candidates known to hitch-hike or associate with mitochondria in axons *in vitro*, only *Pink1* and *Bcl2l2*^28^ were found enriched in ON axonal mitochondria, but were non-significant or de-enriched in SN mitochondria (Fig. 1e-f). This again supports the idea that mitochondrial RNA association is highly dependent on the neuronal cell type.

**Fig. 1.**
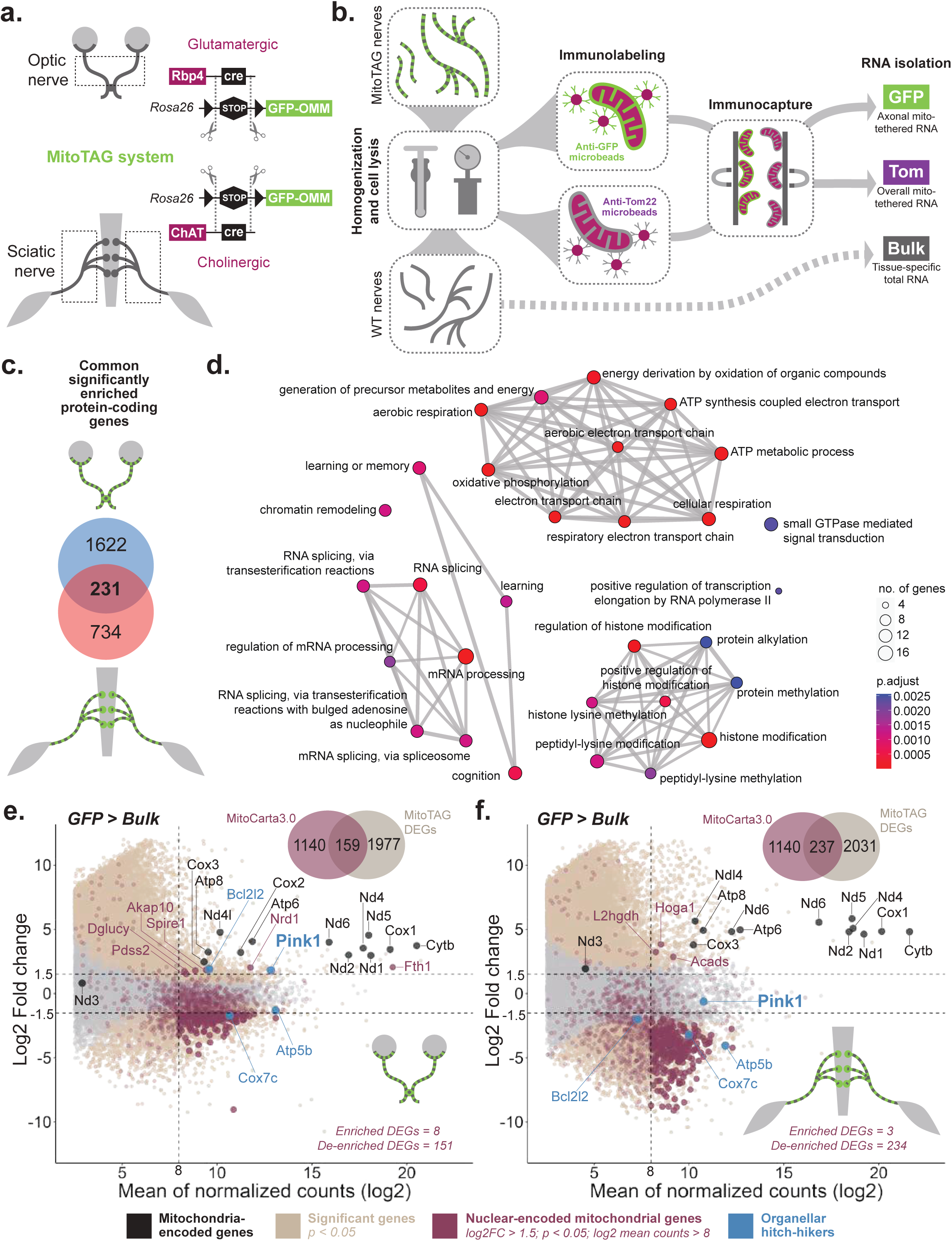
Landscape of mitochondria-mRNA tethering in neurons. **a,** Schematic of the tissues of interest (optic nerve, ON and sciatic nerve, SN) expressing GFP-tagged mitochondria in glutamatergic (Rbp4cre-driven) and cholinergic (ChATcre-driven) axons of ON and SN, respectively, with the help of the MitoTag system. **b,** Schematic of the mitochondria isolation protocol and the groups of interest (GFP, Tom, Bulk). Left to right: nerves homogenized and cells cavitated, mitochondria immunolabelled using microbeads coated with antibodies, immunocaptured under high-field magnet, mitochondria isolated, and total RNA extracted. **c**, Venn diagram showing the number of significantly enriched protein-coding genes that are common and unique from differential gene expression analysis (n = 4) with log2 fold change (log2 FC) > 1.5, Benjamini-Hochberg (adjusted p) p < 0.05, and log2 average gene expression across the groups of comparison (log2 mean counts) > 8 for both ON and SN. **d,** GO enrichment map of the common 231 genes (adjusted p < 0.01) showing its associated top 30 terms after their semantic similarity-based removal (cutoff = 0.9) of the redundant terms. Nodes represent the GO terms connected by edges that represent its semantic similarity, size of dots represent the number of genes, and the colors of the nodes represent the adjusted p-values. **e,f,** MA-plot showing the log2 FC of genes on GFP compared to Bulk versus the log2 mean counts for ON (**e**) and SN (**f**). Differentially expressed genes (DEGs) with adjusted p < 0.05 are colored in beige and p > 0.05 are colored in grey. Mitochondria-encoded genes are colored and labelled in black. Mitocarta3.0 genes are colored in deep ruby and the genes with log2 FC > 1.5 or < −1.5, adjusted p < 0.05, and log2 mean counts > 8 (numbers are shown in the Venn diagram at the top right corner for each tissue) are in the same color with less transparency and bigger size, and the genes with log2 FC > 1.5 are additionally labelled in the same color. Genes that are colored and labelled in blue are previously known organellar hitch-hikers.

### SYNJ2a expression correlates with *Pink1* mRNA localization

To follow up on the cell type specificity observed for the association of the *Pink1* transcript, we confirmed in our dataset that indeed its abundance increased with increasing purification of axonal mitochondria in the ON, but not in the SN (Fig. 2a). The *Pink1* transcript was abundantly detected in ON axons using RNAscope-based *in situ* hybridization in horizontal nerve sections, yet only rarely present in SN axons (Fig. 2b-c).

**Fig. 2.**
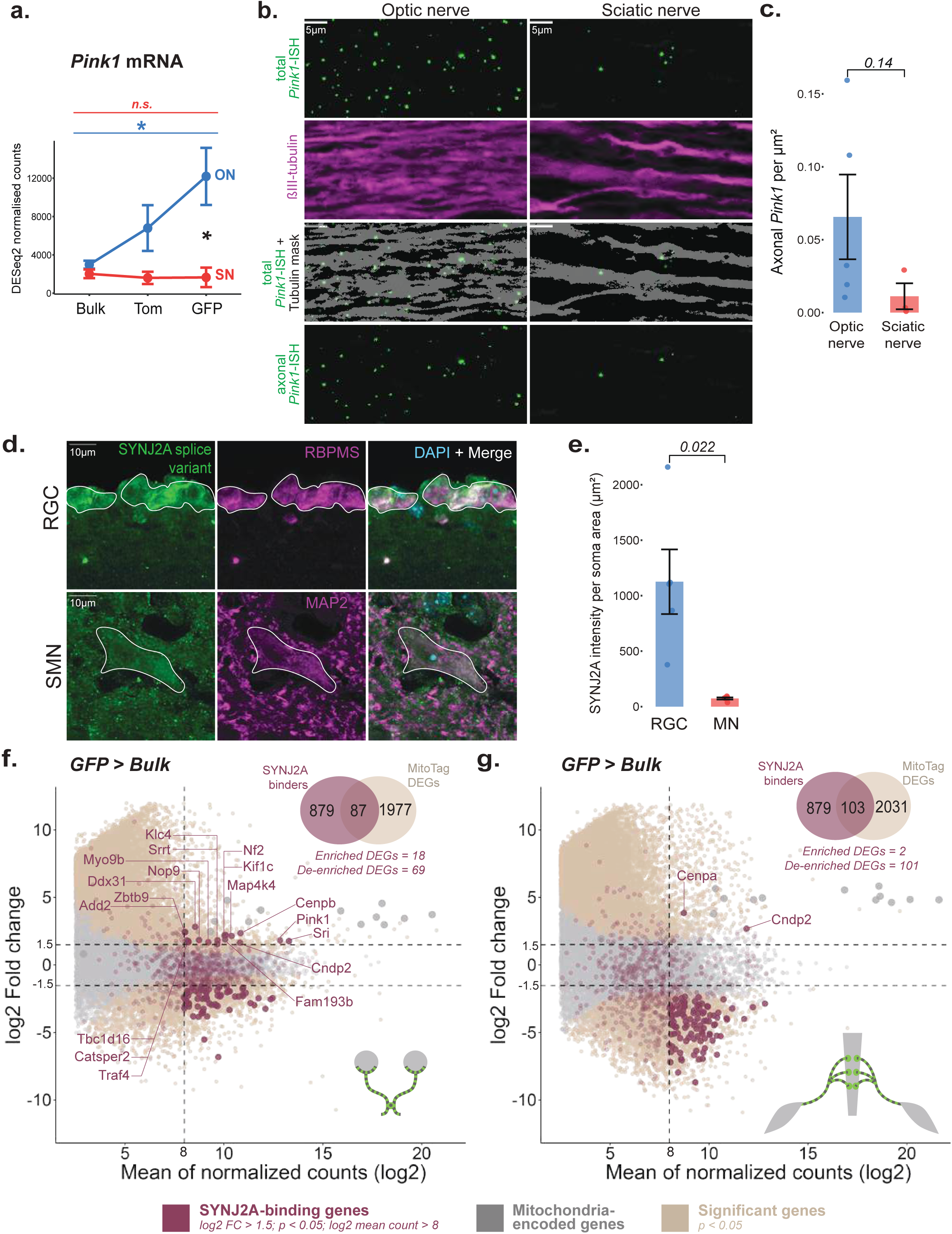
*Pink1* mRNA axonal presence is cell type specific. **a**, Line plot showing the DESeq2-normalised counts of *Pink1* mRNA across GFP, Tom, and Bulk RNA isolated from ON (blue) and SN (red). Two-way ANOVA followed by Tukey’s HSD post-hoc test performed between groups (blue or red) and tissues of interest (black) are depicted (n = 4 pairs of tissues per group, mean ± SEM, *p < 0.005, n.s. not significant). **b**, *In situ* hybridization of *Pink1* mRNA performed on horizontal cryosections of ON (left) and SN (right). Top panel (green) represents total individual endogenous *Pink1* mRNA puncta followed by the axonal marker, ß3-Tubulin (magenta), which is used as a mask to extract the axonal *Pink1* puncta (third panel, mask in grey, mRNA in green). The bottom panel represents axon-only *Pink1*. **c**, Quantification of images as in (**b**) showing the number of axonal *Pink1* transcripts per µm^2^ (n = 3-5 tissue sections, two-tailed Student’s t-test, mean ± SEM) across ON (blue) and SN (red). **d**, Representative images of immunostainings for SYNJ2a using a splice variant specific antibody (green) performed on cross-sections of retina and spinal cord. RBPMS and MAP2 protein (magenta) serve as markers for retinal ganglion cells (RGC) and motor neurons (MN), respectively. **e**, Quantification of the somatic fluorescence intensity of SYNJ2a normalized to the somatic area of the respective neuron (n = 5 tissue sections, two-tailed Student’s t-test, mean ± SEM) for RGC (blue) and SMN (red). **f,g**, MA plots showing log2 FC versus log2 mean counts for ON (**f**) and SN (**g**). Genes that are differentially expressed on GFP compared to Bulk with Benjamini-Hochberg-adjusted p < 0.05 are colored in beige and p > 0.05 are colored in grey. Mitochondria-encoded genes are colored in dark grey. SYNJ2a-binding mRNAs are colored in deep ruby, the genes with log2 FC > 1.5 or < −1.5, adjusted p < 0.05, and log2 mean counts > 8 (numbers are shown in the Venn diagram at the top right corner for each tissue) are in the same color with less transparency and bigger size, and the genes with log2 FC > 1.5 are additionally labelled in the same color. Scale bars: 10µm (**b**), 5µm (**d**).

As the expression of SYNJ2a is a prerequisite for the mitochondrial association of the *Pink1* mRNA^21^, we hypothesized that the expression of the mitochondria-specific isoform of this RBP could underlie some of the cell type diversity. Using an isoform-specific antibody, we readily detected expression of SYNJ2a in retinal ganglion cells (RGCs), which give rise to the axons in the ON (Fig. 2d-e). On the contrary, expression of SYNJ2a in the somas of lumbar motor neurons (MN) was barely detectable (Fig. 2d-e). Thus, lumbar MNs lack the ability to transport the *Pink1* mRNA via the mitochondrial mRNA anchoring complex, explaining its absence from axonal mitochondria derived from their axons.

Given the results for *Pink1*, we tested whether the differential expression of SYNJ2a could explain more of the cell type-specific differences between ON and SN axonal mitochondria. We have previously sequenced all transcripts bound by SYNJ2a in HEK293 cells^21^. Upon comparison of this dataset with the transcripts significantly enriched or de-enriched on axonal mitochondria in SN and ON tissue, we confirmed that SYNJ2 transcripts were more frequently enriched in ON axonal mitochondria compared to the SN. In total, 18 SYNJ2a-binding transcripts reached statistical significance for enrichment on ON axonal mitochondria, whereas only 2 transcripts coincided with the mRNAs associated with SN axonal mitochondria (Fig. 2f-g). This suggests that indeed the differential expression of the RBP SYNJ2a underlies some of the cell type specific differences in the landscape of transcripts associated with axonal mitochondria.

Also other RBPs have been suggested to locate transcripts to mitochondria. Of these, datasets of transcripts bound to the RBPs CLUH and SYNJ2BP have been published^29,30^. However, neither of these RBP-bound transcripts showed a similar preference for any of the mRNAs associated with SN or ON axonal mitochondria (Fig. S2a-d). Likewise, transcripts hitching a ride on endolysosomes via BORC-mediated transport^5^ did not show a preferential enrichment in our datasets (Fig. S2e-f). This suggests that mitochondria-associated mRNAs make up a specific subpopulation of axonal transcripts, and that the selective expression of SYNJ2a is responsible for the ability of RGCs to transport *Pink1* mRNA and other SYNJ2a substrates.

### *Mapt* mRNA transport depends on the mitochondrial mRNA anchoring complex

As nuclear-encoded mitochondrial genes were underrepresented in the mRNAs associated with axonal mitochondria in the optic nerve, we performed gene ontology (GO) analysis to discover whether another category was enriched on ON axonal mitochondria. Intriguingly, next to GO terms related to mitochondria, mainly driven by the mtDNA-endcoded genes, cellular components connected to cytoskeletal elements were highly enriched. Among those, “microtubule” was the most enriched GO term (Fig. 3a). Focusing on microtubule-associated proteins (MAPs), we observed that the enrichment of *Mapt*, *Map7* and *Map4* was again highly ON specific (Fig. 3b), with especially *Mapt* even being de-enriched in the SN axonal mitochondria (Fig. 3c). This behavior was highly reminiscent of the distribution of *Pink1*, as its abundance increased with increasing purification of axonal mitochondria in the ON, but not in the SN (Fig. 3d). The *Mapt* transcript was abundantly detected in ON axons using RNAscope-based *in situ* hybridization in horizontal nerve sections, as was its protein product Tau (Fig. 3e-h). In contrast, less *Mapt* mRNA and Tau immunostaining was detectable in SN horizontal nerve sections (Fig. 3e-h). Importantly, we used a probe that detects all *Mapt* isoforms, including the “Big-Tau” isoform containing the exon 4a that has been reported to occur specifically in peripheral MNs^19^. Given that peripheral MNs express a different isoform of Tau, the need to localize the *Mapt* transcript for local translation of Tau may only apply to the centrally expressed isoforms.

**Fig. 3.**
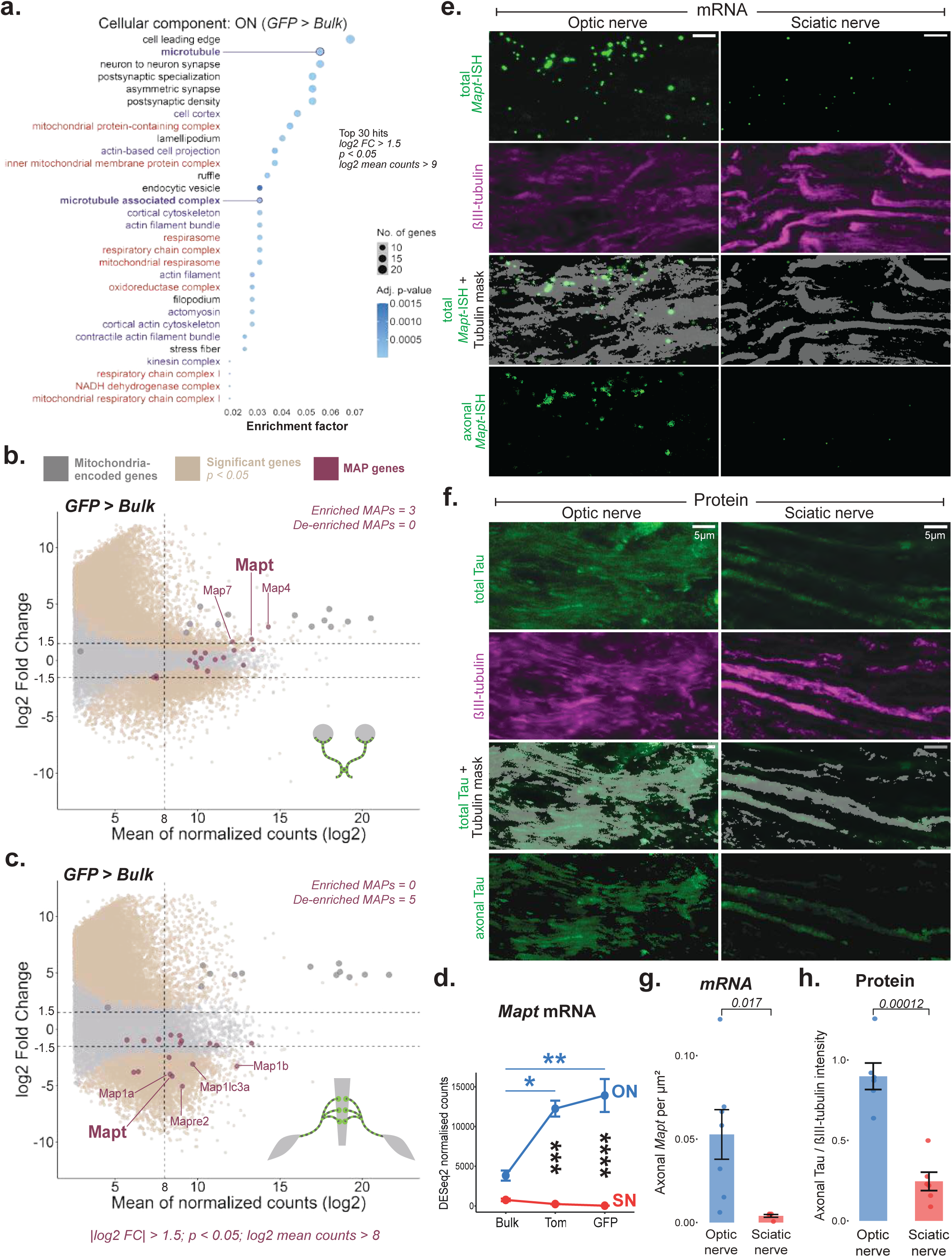
*Mapt* mRNA associates with ON mitochondria. **a**, Dot plot showing the enrichment factor of the top 30 GO cellular component terms with the least q-values performed on the significantly enriched genes of GFP compared to Bulk for ON (log2 FC > 1.5, adjusted p < 0.05, log2 mean counts > 9). Size of the dots represent the number of genes present within the term and the adjusted p values are represented in shades of blue. Enrichment was assessed using Fischer’s exact test followed by Benjamini-Hochberg correction. Terms related to microtubule/cytoskeleton are colored in purple while terms related to mitochondria are colored in red. **b,c,** MA plots showing log2 FC versus log2 mean counts for ON (**b**) and SN (**c**). Genes that are differentially expressed on GFP compared to Bulk with Benjamini-Hochberg-adjusted p < 0.05 are colored in beige and p > 0.05 are colored in grey. Mitochondria-encoded genes are colored in dark grey. Microtubule-associated protein (MAP) genes are colored in deep ruby and the genes that surpass the threshold of log2 FC > 1.5, adjusted p < 0.05, and log2 mean counts > 8 are additionally labelled in the same color. **d,** Line plot showing the DESeq2-normalised counts of *Mapt* across GFP, Tom, and Bulk isolated from ON (blue) and SN (red). Two-way ANOVA followed by Tukey’s HSD post-hoc test performed between groups (blue or red) and tissues of interest (black) are depicted (n = 4 pairs of tissues per group, mean ± SEM, *p < 0.001, **p < 0.0001, ***p < 0.00001, ****p < 0.000001). **e,f,** RNAseq result of *Mapt* is validated at the mRNA (e) and protein (f) levels on horizontal sections of ON (left) and SN (right). Top panel (green) represents total individual endogenous *Mapt* mRNA puncta (**e**) and total Tau level (**f**) followed by the axonal marker, ß3-Tubulin (magenta), which is used as a mask to extract the (third panel, mask in grey, mRNA/protein in green) axonal *Mapt* puncta (**e**) and axonal Tau levels (**f**). The last bottom panel axon-only *Mapt* (**e**) and Tau (**f**) signal. **g,h,** Quantification of number of axonal *Mapt* puncta normalized per µm^2^ (e,g; n = 5-7 tissue sections, mean ± SEM, two-sided Student’s t-test) and axonal mean Tau fluorescence intensity normalized to that of ß3-Tubulin (**f,h**; n = 6 tissue sections, mean ± SEM, two-tailed Student’s t-test) across ON (blue) and SN (red). Scale bars: 5µm (**e,f**)

### Loss of mitochondrial hitch hiking impairs Tau sorting and axon growth

The similarity to *Pink1* mRNA distribution prompted us to test whether the mitochondrial association of *Mapt* mRNA would also be detectable in cultured hippocampal neurons. Indeed, we detect association of the *Mapt* transcript with mitochondria in both soma and neurites, and this association is sensitive to knock down of SYNJ2BP (Fig. 4a-c). The signal was specific, as we observed no signal for the negative control probe targeting the bacterial transcript *DapB* as well as no change in the mitochondrial association of the control transcript *Ppib* (Fig. S4a-b). We thus propose that *Mapt* is a substrate for mitochondrial hitch-hiking via the SYNJ2BP/SYNJ2a complex in glutamatergic neurons. In contrast to *Pink1* mRNA^21^, however, also the overall abundance of the *Mapt* transcript decreased (Fig. 4d), suggesting that non-mitochondrial *Mapt* mRNA may be actively degraded.

**Fig. 4.**
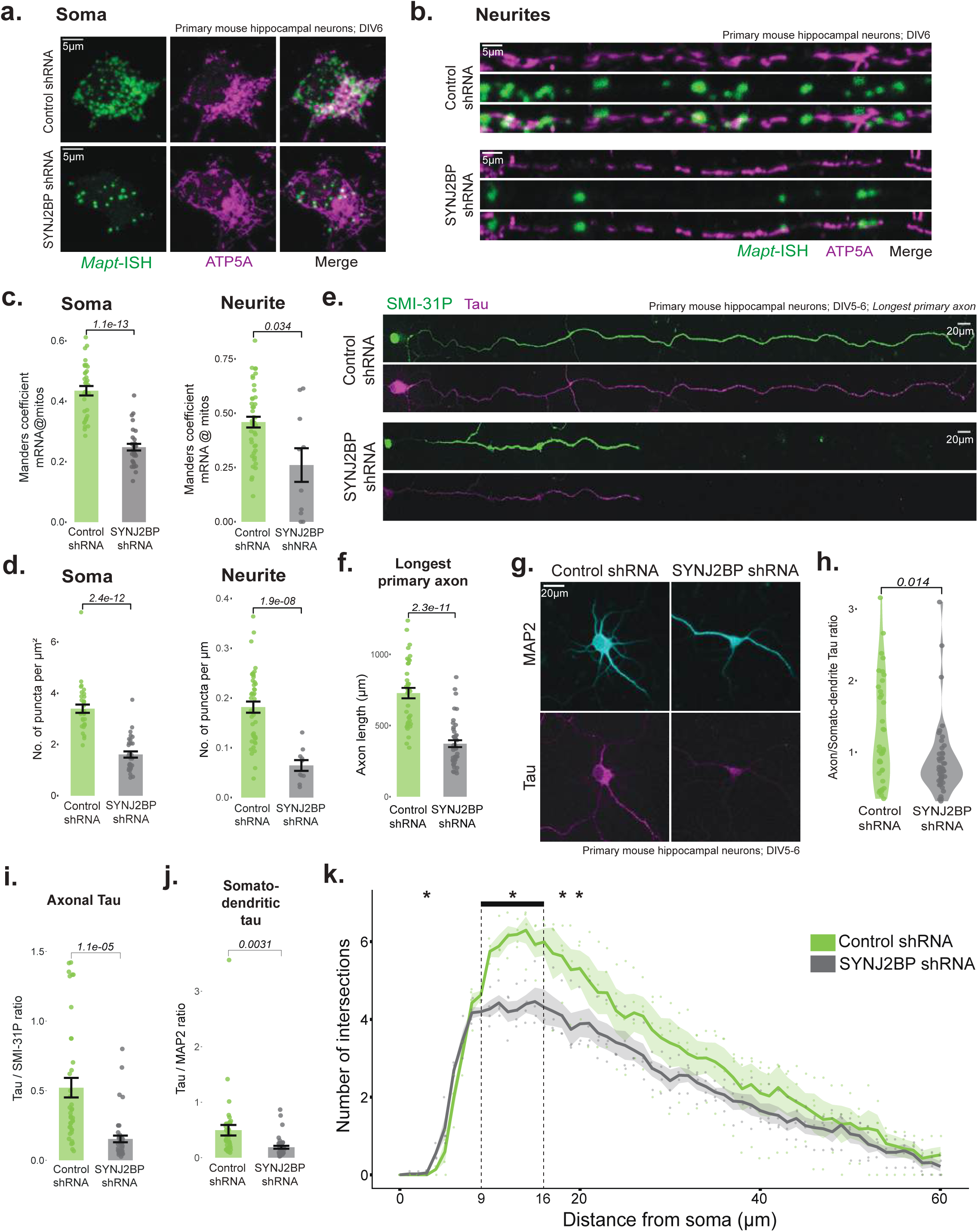
Loss of *Mapt*-mitochondria association affects Tau levels and neurite growth. **a,b,** Representative images showing colocalization of endogenous *Mapt* (green; left) puncta with Atp5α-stained mitochondria (magenta; middle) after 5 days of control or SYNJ2BP knockdown in cultured hippocampal neurons (DIV6). Soma (**a**) and digitally straightened neurites (**b**). White color appearing on the “Merge” (right) panel visually represents the colocalization. **c,d,** Mander’s correlation coefficient of *Mapt* puncta on mitochondria (**c**) and the number of puncta (**d**) normalized per µm^2^ (right) area of soma (n = 31-33) and per µm length of neurite (n = 10-43) in control or SYNJ2BP knockdown neurons. **e** Immunostaining images of axonal marker, Smi31p (green), and Tau (magenta) after 4 days of control or SYNJ2BP knockdown (DIV5) **f** Quantification of the longest axon length in images as in (**e**) (n = 38-44 axons). **g**, Immunostaining of somato-dendritic marker, Map2 (cyan), and Tau (magenta) after 5 days of control or SYNJ2BP knockdown (DIV6). **h**, Violin plot showing the ratio of axonal to somato-dendritic Tau fluorescence intensity of neurons as in (**g**) (n = 38-44 neurons). **i**, Quantification of the ratio of axonal Tau fluorescence intensity normalized to Smi31p of neurons as in (**g**) (n = 38-44 neurons) **j,** Quantification of the ratio of somato-dendritic Tau fluorescence intensity normalized to Map2 of neurons as in (**g**) (n = 38-44 neurons). **k**, Line plot of the distance from soma as a function of the number of neurite intersections after 5-6 days after knockdown (n = 4 biological replicates, mean ± SEM). Two-way ANOVA is done between the knockdown conditions (green or dark grey) and the step radii (step size = 1µm, 60 steps in total) followed by Tukey’s HSD post-hoc test for multiple comparisons. Continuous statistical significance (*p < 0.05) is shown between 9-16µm, and discontinuously observed at radii, 4, 18, and 20µm from the center of soma. All of the aforementioned experiments are performed on cultured primary mouse hippocampal neurons transduced with either control (green) or SYNJ2BP (dark grey) shRNA. Mean ± SEM are shown (except h) and the p values are calculated using two-tailed Student’s t-test, unless otherwise mentioned. Scale bars: 5µm (**a,b**), 20µm (**e,g**)

To analyze the functional impact of mitochondrial hitch-hiking of *Mapt*, we analyzed the Tau levels upon knock down of SYNJ2BP during neuronal differentiation *in vitro*. Strikingly, knock down of SYNJ2BP already visibly decreased the length of the axonal extension on day *in vitro* (DIV) 5 (Fig. 4e-f and S4c). Tau levels were also decreasing in the somatic area, as expected from a loss of *Mapt* mRNA abundance (compare Fig. 4a-d), yet the difference became even more striking in the axon (Fig. 4g-j). Thus, loss of mitochondrial association of *Mapt* leads to decreased expression and mis-sorting of Tau during neurodevelopment *in vitro*.

Given our observation that several cytoskeletal mRNAs are associating with axonal mitochondria, we reasoned that the effect of SYNJ2BP on neurite growth may not be restricted to axons. Indeed, Sholl analysis^31^ revealed that while the overall ramification of SYNJ2BP knock down neurons was similar to control shRNA treated neurons, the number of intersections decreased significantly between 9-16 µm away from the soma of SYNJ2BP knock down neurons. Thus, mitochondrial hitch-hiking of mRNAs, including *Mapt,* sculpts the local cytoskeleton to sustain neurite outgrowth.

## Discussion

Our results show that there are fundamental differences between the cholinergic PNS and glutamatergic CNS axons with regards to their repertoire of mitochondrially associated mRNAs. We only find that a select number of nuclear-encoded mitochondrial genes attach to mitochondria (Fig. 1e-f and S1c-d). In ON derived axonal mitochondria compared to their expression levels in bulk tissue (Fig. 1e-f), this includes the transcript for *Pink1*, which we had previously shown to be transported in RGC axons. Similarly, another transcript previously found associated with neuronal mitochondria, *Bcl2l2*^28^ is significantly associated with ON axonal and bulk mitochondria (Fig. 1e-f and S1c-d). Other candidates previously shown to associate with neuronal mitochondria, *Cox7c*^22^ and *Atp5*^32^, are not enriched. As *Cox7c* is thought to associate via co-translational import of its encoded protein, its interaction may be too labile to survive the immuno-isolation protocol. Likewise, CLUH is responsible for co-translational attachment of mRNAs to mitochondria^29,33,34^, yet we do not find evidence for an enrichment of CLUH targets in ON or SN axonal mitochondria (Fig. S2a-b). Thus, our dataset only includes transcripts whose association is stabilized beyond co-translational targeting.

The cell type specific differences in the mitochondrial RNA association landscape can be correlated to the differential expression of the RBP SYNJ2a in these cell types. We show that the expression of the SYNJ2a isoform is lower in lumbar MNs (Fig. 2e), while RGCs express comparatively higher levels, fitting to the enrichment of SYNJ2a-bound transcripts associating with ON axonal mitochondria (Fig. 2f). Likewise, the *Mapt* mRNA follows this pattern and was thus confirmed *in vitro* as a further substrate of the mitochondrial hitch-hiking complex (Fig. 4a-c). This implies that MNs lack mitochondrial hitch-hiking of *Mapt* and *Pink1*. Thus, MNs need to have evolved alternative strategies to sustain mitochondrial quality control and microtubule stability in axons. For *Pink1*, it is well documented that other mitophagy pathways can compensate for its absence^35,36^. PINK1-independent mitophagy has indeed been shown to be responsible for basal mitophagy in MNs^37^, yet whether this is true for damaged-induced mitophagy remains to be tested. For Tau, the difference may be explained by the selective expression of the “Big Tau” splice variant in the PNS^19^. Not much is known about the differences between this variant and its CNS enriched versions. We propose that CNS Tau may need to be replenished more frequently and thus relies on a higher presence of *Mapt* mRNA in axons (Fig. 3e,g). In contrast, Big Tau may have higher stability and thus translation in the soma and protein-based sorting mechanisms, or transport via hnRNP R-containing ribonuclear particles^14^, may suffice to supply Tau to MN axons.

Our data supports the idea that trafficking of the *Mapt* mRNA via mitochondrial hitch-hiking is necessary to maintain Tau levels in glutamatergic CNS neurons. At the level of the mRNA, we now have revealed that SYNJ2BP is necessary for the mitochondrial association of the *Mapt* mRNA. Knock down of SYNJ2BP reduced the overall levels of *Mapt* and Tau, with its biggest impact on the axonal levels of Tau, suggesting a role of this mitochondrial protein in *Mapt* transport, similar to the mitochondrial hitch-hiking of *Pink1*^21^. While we do not formally show a dependence on SYNJ2a, it is highly likely that *Mapt* requires both SYNJ2BP and SYNJ2a, due to its cell type specific mitochondrial association (Fig. 3b-c) which is only seen for SYNJ2 but not SYNJ2BP-dependent transcripts (Fig. 2e-f and S2c-d). Interestingly, mitochondrial hitch-hiking occurs in a translationally repressed state^38^, thus there need to be axon-specific mechanisms to activate the localized translation of Tau. This has been suggested to depend on an oligopyrimidine tract within its 5’-UTR, which drives axonal protein translation in response to mTORC1 activity^15^. Given that mTORC1 activation occurs at the endolysosomal surface^39^ and that axonal translation occurs at contact sites between mitochondria and endolysosomes^20,40^, this suggests that the formation of such organellar contact sites may be an important driver of localized Tau production. Importantly, while we use *Mapt* to show the importance of mitochondrial mRNA association for transport and local translation of Tau, this applies to all mitochondrially associated mRNAs, which are highly enriched in cytoskeletal elements in the ON axons. Thus, loss of the mitochondrial hitch-hiking complex not only stunts axonal growth *in vitro* (Fig. 4e-f), but also affects dendritic arborization (Fig. 4k). This matches well with previously established roles for mitochondria to determine axonal and dendritic branch sites^7,41,42^, which among other functions depends on the associated local translation of cytoskeletal mRNAs^7^.

It is interesting to note that this potential difference in Tau homeostasis reflects the susceptibility of neurons to form Tau tangles, which dominantly observed in the CNS but not in the PNS in the context of AD and related tauopathies. Loss of mitochondrial motility has been observed in aging^43^ and in AD mouse models^44,45^ and our data now suggest that this may lead to a vicious cycle as reduced mitochondrial motility will lead to a drop in *Mapt* availability to replenish the axonal pool of Tau. This could contribute to the mislocalization of Tau protein in disease and promote the formation of Tau Tangles^12^. This in turn further reduces mitochondrial motility^46,47^. While this model still needs to be tested in the disease context, our data reveal a fundamental relationship between mitochondrial mRNA hitch-hiking and Tau homeostasis in neurons.

## Supplementary figures

**Fig. S1.**
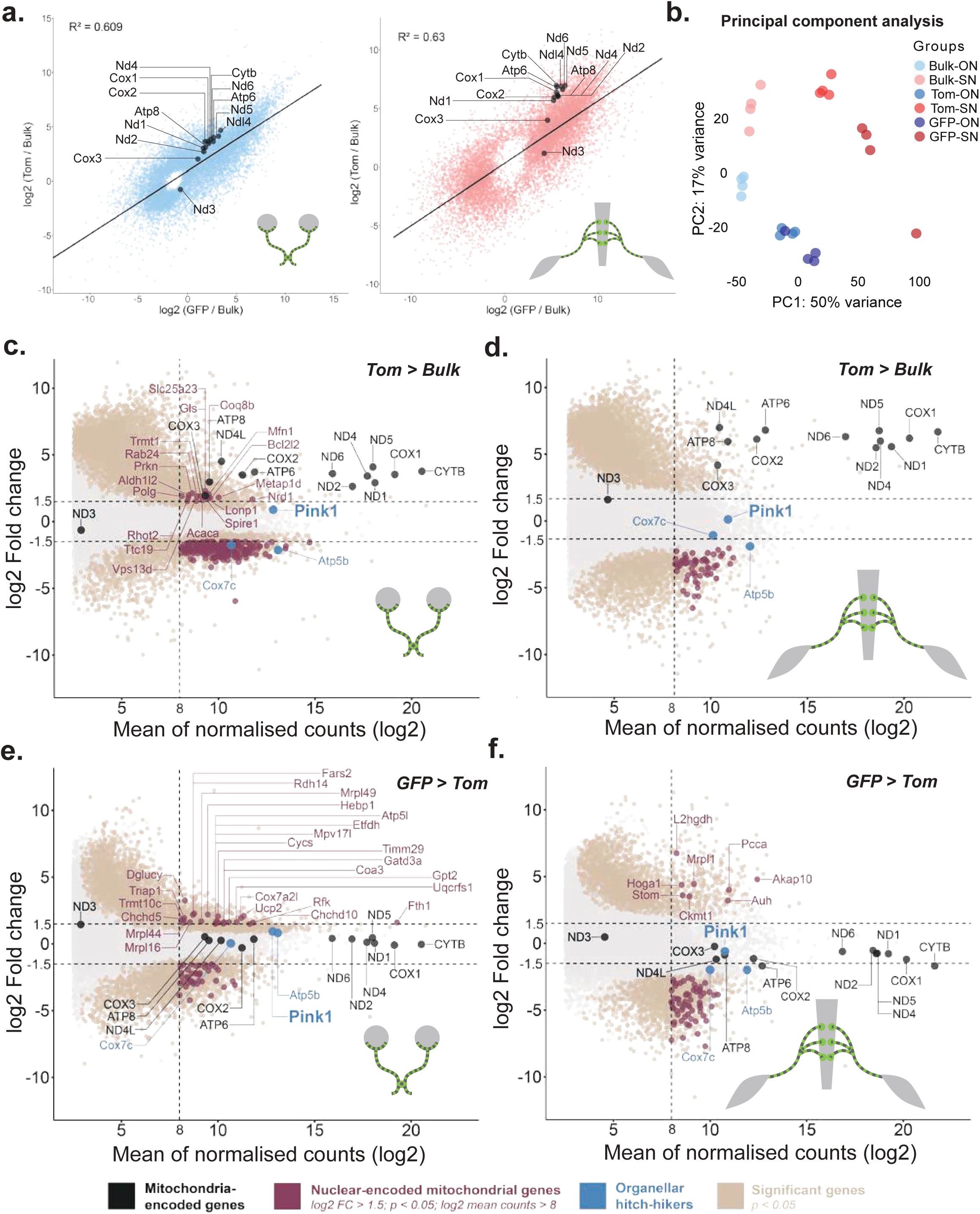
Differential gene expression analysis of isolated mitochondria. **a**, Scatter plot shows the mean DESeq2-normalized counts of GFP versus Tom that is further normalized to Bulk (log2GFP/Bulk and log2Tom/Bulk) for ON (light blue) and SN (light red). These DEGs pass Benjamini-Hochberg-adjusted p < 0.05 in differential gene expression analysis (n = 4 pairs of tissues). Linear regression line and the coefficient of determination (top left corner of each graph) are depicted. Mitochondria-encoded genes are colored and labelled in black. **b**, Principle component analysis (PCA) of variance-stabilized transformed (vst) counts. Lighter to darker shades of blue and red represent ON and SN, respectively. Each dot depicts one biological replicate. Percentage variance explained by each principle component is described on the axes. **c-f**, MA-plot showing the log2 fold change (log2 FC) of genes on Tom compared to Bulk versus the log2 average gene expression across (log2 mean counts) Tom and Bulk for ON (**c**) and SN (**d**), and genes on GFP compared to Tom for ON (**e**) and SN (**f**). Differentially expressed genes with Benjamini-Hochberg-adjusted p < 0.05 are colored in beige and p > 0.05 are colored in light grey. Mitochondria-encoded genes are colored and labelled in black. Mitocarta3.0 genes with log2 FC > 1.5 or < −1.5, adjusted p < 0.05, and log2 mean counts > 8 are colored in deep ruby and the genes with log2 FC > 1.5 are additionally labelled in the same color. Genes that are colored and labelled in blue are previously known organellar hitch-hikers.

**Fig. S2.**
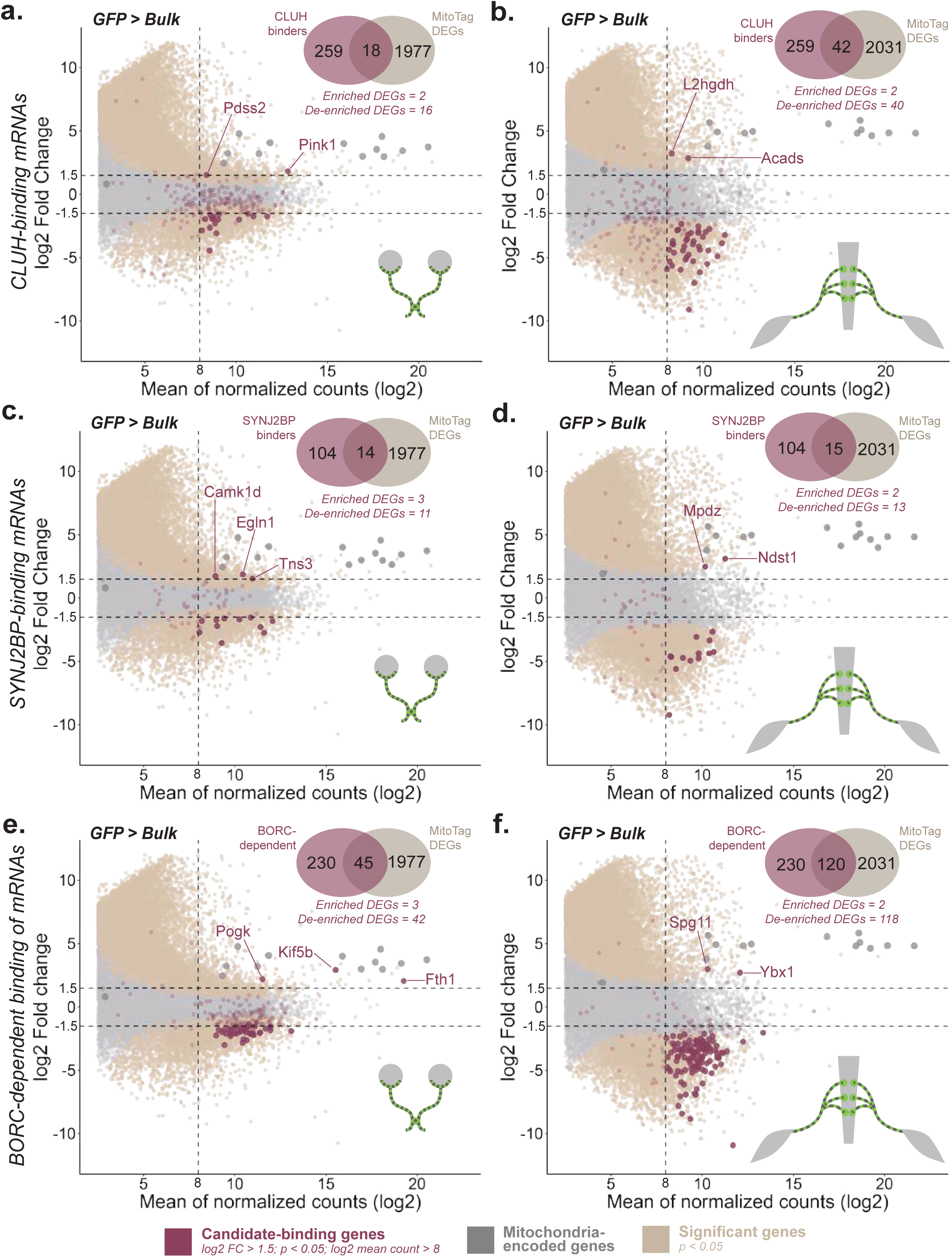
Axonal mitochondria-enrichment of RBP-bound mRNAs. **a-f**, MA-plot showing the log2 fold change (log2 FC) of genes on GFP compared to Bulk versus the log2 average gene expression across (log2 mean counts) GFP and Bulk for ON (**a,c,e**) and SN (**b,d,f**). DEGs with Benjamini-Hochberg-adjusted p < 0.05 are colored in beige and p > 0.05 are colored in grey. Mitochondria-encoded genes are colored in dark grey. Gene sets that are previously identified to interact with protein complexes are colored in deep ruby and the genes with log2 FC > 1.5 or < - 1.5, adjusted p < 0.05, and log2 mean counts > 8 (numbers are shown in the Venn diagram) are in the same color with less transparency and bigger size, and the genes with log2 FC > 1.5 are additionally labelled in the same color. Gene sets that are known to interact via via BORC-dependent complex (**a,b**), SYNJ2BP (**c,d**), and CLUH (**e,f**) are shown.

**Fig. S3.**
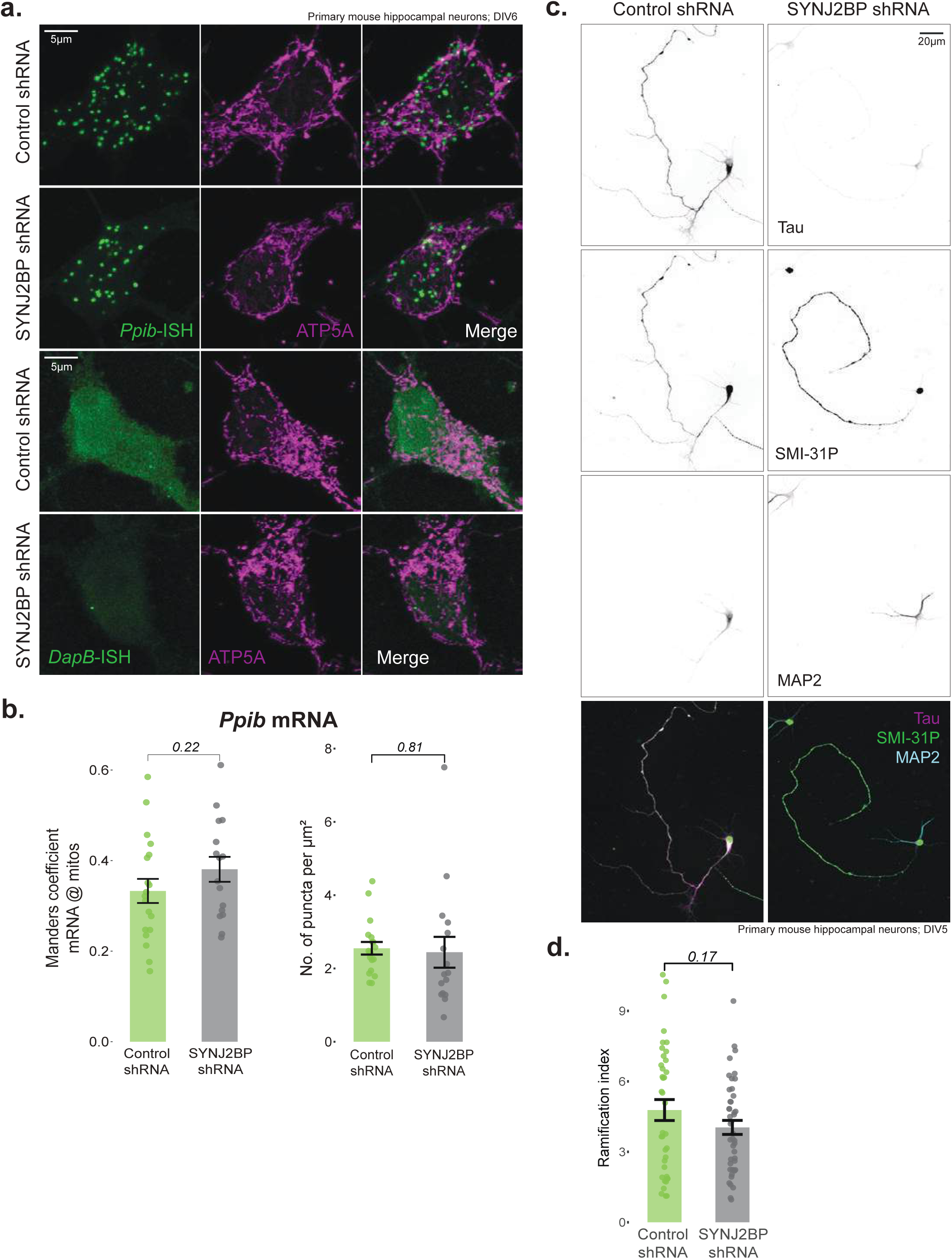
SYNJ2BP knock down causes no general RNA redistribution yet alters axon length. **a**, Representative images of RNAscope (green; left) of a ubiquitously expressed transcript, *Ppib*, and a negative control, *DapB*, immunostained with mitochondrial marker, Atp5α (magenta; middle) after 5 days of knockdown. White color appearing in the “Merge” (right) panel is the visual representation of colocalization. **b**, Mander’s correlation coefficient of images as in (**a**). *Ppib* (left) and the number of puncta normalized per µm^2^ (right) area (n = 16-19 soma). **c**, Immunofluorescence images after 4 days of knockdown, stained for Tau (top panel), Smi31p (axonal marker; middle panel), and Map2 (somato-dendritic marker; bottom panel). **d**, Ramification index quantified by the total number of neurite intersections normalized to the number of primary neurites (n = 38-44 neurons). All of the aforementioned experiments are performed in cultured primary mouse hippocampal neurons transduced with control (green) and SYNJ2BP (dark grey) shRNA on DIV1. Mean ± SEM are shown with p values calculated using two-tailed Student’s t-test, unless otherwise mentioned. Scale bars: 5µm (a), 20µm (c).

## Acknowledgments

We thank J. Lindner for technical support and the members of the Harbauer and Misgeld laboratories for their support and many fruitful discussions. We are grateful for the support of Rinho Kim from the NGS core of the MPI for Biochemistry (RRID:SCR_025746). We are also grateful to R. Kasper/E. Laurell/C. Polisseni and M. Spitaler/M. Oster from the Imaging facilities of the MPIs for Biological Intelligence and Biochemistry, respectively, for assistance with microscopy (RRID:SCR_026797, RRID:SCR_025739).

## Funding

Work in **ABH’s** lab is supported by

Max Planck Society
DFG (DFG, TRR353 – ID 471011418, SPP2453 – ID 541742535)
European Union (ERC StG Project 101077138 — MitoPIP)
Germany’s Excellence Strategy within the framework of the Munich Cluster for Systems Neurology (EXC 2145 SyNergy – ID 390857198)
Schram Foundation (ID T0287/46550/2025)

Work in **TM’s** lab is supported by

German Research Foundation (DFG, Mi 694/8-1 “, FOR Immunostroke (Mi 694/9-1/2 – ID 428663564, TRR 274/1 Projects C02, Z01, respectively – ID 408885537).
German Center for Neurodegenerative Diseases (DZNE).
DFG instrumentation grant (INST95/1755-1 FUGG, ID 518284373).
Chan Zuckerberg Initiative (CZI) grant for Creating Community & Building Capacity.
Germany’s Excellence Strategy within the framework of the Munich Cluster for Systems Neurology (EXC 2145 SyNergy – ID 390857198).

Work in **RF’s** facility is supported by

Germany’s Excellence Strategy within the framework of the Munich Cluster for Systems Neurology (EXC 2145 SyNergy – ID 390857198).
German Center for Neurodegenerative Diseases (DZNE).
Helmholtz Center Munich, German Research Center for Environmental Health GmbH, Neuherberg, Germany.

## Author contributions

Conceptualization: ABH

Methodology: ABH, HMM, AER, AMP, MB, IMCG, MB, LRA, RF

Investigation: HMM, AER

Visualization: HMM

Supervision: ABH, TM

Writing – original draft: ABH, HMM

Writing – review & editing: all authors

## Data and materials availability

Requests for resources and reagents should be directed to and will be fulfilled by the lead contact Angelika B. Harbauer. Microscopy data reported in this paper will be shared by the lead contact upon request. Any additional information required to reanalyze the data reported in this paper is available from the lead contact upon request. Sequencing data will be made available upon publication in a publicly available depository.

## Methods

### Animals

All animal experiments were conducted during their light phase and all animals had *ad libitum* access to food and water. Only males with 12-22 months of age were used for transcriptomics. Wildtype (C57BL/6 N background) males and females between 10-20 months of age were used for RNAscope and immunohistochemistry. All mouse experiments were approved by the Max Planck Institute for Biological Intelligence/TUM and were adhering to the regulations of the Government of upper Bavaria. Female Wistar rats aged 4 to 7 months were used for immunization in accordance with the German Animal Welfare Law and with the approval of the local authorities of Upper Bavaria, Germany (ROB-55.2Vet-2532.Vet_03-22-25).

#### MitoTag

These animals were housed and maintained at the Technical University of Munich, within the Institute of Neuronal Cell Biology. A 12:12 h light-dark cycle was maintained for MitoTag animals. To generate cell-type-specific GFP-tagged mitochondria, the reporter strain, MitoTag*^1^* (GFP-OMM^fl/fl^; Jackson Labs, 032675) was crossed with the following Cre-driver lines: Rbp4-Cre^48^ (031125-UCD; Mutant Mouse Resource and Research Center, MMRRC), ChAT-Cre^49^ (Jackson Labs, 006410).

#### Wildtype

Wildtype animals were housed and maintained in Max Planck Institute for Biological Intelligence Martinsried with 14:10h light-dark cycle.

### RNA extraction from bulk and mitochondrial pellets

MitoTag-positive and wildtype animals were euthanized by either isoflurane or carbon-di-oxide overdose, respectively, followed by PBS transcardial perfusion for MitoTag animals. Fresh tissues of interest (kept on ice throughout the procedure) were immediately processed for performing immunocapture of mitochondria using the protocol adapted from Fecher et al. 2019. Briefly, the tissues were transferred to a dounce tissue homogenizer (DWK Life Sciences; 357542) containing 2mL of isolation buffer (IB), made using (in mM): 220 mannitol, 80 sucrose, 10 HEPES, 1 EDTA, set to pH 7.40 using 1M KOH, and freshly supplemented with 1% fatty acid-free bovine serum albumin (BSA; Sigma Aldrich; A2153), and 1X cOmplete protease inhibitor (Roche; 04 693 132 001). Using a type-A (loose) pestle, the tissues were homogenized with 10-15 strokes. Further, the cells were opened using the nitrogen cavitation system (Parr Instrument Company; model 4635–39) inside a cell disruption vessel at 800psi for 10mins. The entire lysed extract (2mL) was filtered through a 30µm separation filter (Miltenyi Biotec; 130-041-407) into the immunoprecipitation buffer (IP), made using (in mM): 137 KCl, 2.5 MgCl_2_, 3 KH_2_PO_4_,10 HEPES, 1 EDTA, pH 7.40, and freshly supplemented with 1% fatty acid-free BSA and 1X cOmplete protease inhibitor. The ratio of IB:IP was 1:5. For immunocapture of axonal mitochondria and overall mitochondria, 25µL of microbeads coated with antibodies against GFP (Miltenyi Biotec; 130-091-288) and Tom22 (Miltenyi Biotec; 130-127-693) were used. The microbeads were added to the IB lysate-IP mixture and incubated at 4-7°C for 90 mins with gentle rotation. To separate the microbead-coated mitochondria, the LS columns (Miltenyi Biotec; 130-042-401) were equilibrated with 3mL of IP after placing it in the MidiMACS separator (Miltenyi Biotec; 130-042-302). After letting the mixture to completely flow through, the columns were removed from the separator, 2mL of IP was added and the mitochondria were mechanically isolated using the plunger. The mitochondria were further pelleted at 10,000xg for 12 mins at 4-7°C. The pellet was immediately snap frozen in liquid nitrogen and later total RNA was extracted with the help of RNeasy Mini kit (Qiagen; 74104), resulting in axonal (GFP) and overall mitochondria (Tom22)-tethered RNA.

For phenol-chloroform-based RNA extraction from bulk tissues, fresh ON and SN tissues were transferred to the dounce tissue homogenizer containing 1mL TRI reagent (Sigma Aldrich; T9424). The tissues were homogenized and lysed using a type-B pestle (tight) with 30-33 strokes. To remove debris, the mixture was centrifuged at 12,000xg for 10 mins at 7°C. The supernatant was mixed with 0.2mL chloroform and the mixture was vigorously shaken for 15 seconds and allowed to stand for 7 mins at room temperature. Next, the mixture was centrifuged at 12,000xg for 15mins at 7°C, phase-separating RNA into a clear aqueous top layer, which was transferred to the RNeasy Qiagen kit for further RNA extraction, resulting in the tissue-specific RNA (Bulk). Total RNA for all the groups of interest was eluted in 23-25µL of RNAse-free water and stored at −80°C until transcriptomics.

### Bulk transcriptomics

RNA concentration was measured using Qubit Flexi-Fluorometer (Q33327). NEB Next Single Cell/Low Input RNA Library Prep Kit (New England Biolabs; E6420) was used for cDNA library preparation after enriching mRNA using Poly oligo-dT primers. The libraries were quantified using 4200 TapeStation System (G2991BA, Agilent) and sequenced using Illumina NovaSeq 6000 (2x 60bp paired-end sequencing) at a depth of 25 million on average.

Raw data were trimmed using Trim Galore^50^, which is a wrapper script around Cutadapt^51^ and FastQC^52^ to remove Illumina adapters and performing data quality control steps, respectively. Trimmed paired-end data were mapped to *Mus musculus* GRCm38.p4 (mm10) genome sequence using HISAT2^53^ using the UCSC mm10 genome build. Samtools^54^ was used to convert the resulting mapped SAM files to BAM files. featureCounts^55^ was used to count the reads from the converted BAM files with the help of UCSC GTF reference. Rest of the analyses were performed in R Studio 4.2.0 (https://www.R-project.org/). Differential gene expression analysis was performed separately for optic nerve and sciatic nerve using DESeq2^56^ package with GFP > Tom, GFP > Bulk, and Tom > Bulk as contrasts.

Genes with Benjamini-Hochberg-adjusted *p* < 0.05 were considered as significant differentially expressed genes (DEGs). To further narrow down significantly enriched or de-enriched DEGs, a cutoff of log2 fold change (log2 FC) > 1.5 or < −1.5 was applied, respectively. Since low expressed genes with extremely high fold change might not reflect biological meaning, two additional cutoffs were applied to only include the DEGs with log2 FC < 12 and log2 (baseMean + 1) > 8 (reflects overall average expression across the groups of comparison). Significantly enriched genes (log2 FC > 1.5) with log2 (average expression across groups) > 9 that are common between ON and SN were used to create the enrichment map. PCA was performed using the variance-stabilized transformed counts (extracted from DESeq2). Any further downstream analyses, statistical tests, and visualizations were performed in R 4.2.1 using tidyverse (ggplot2, tibble, and tidyr), gprofiler2, and clusterProfiler.

### *In-situ* hybridization and immunofluorescence

#### Cultured cells

Primary mouse hippocampal neurons were isolated, maintained, and transduced with lentiviral particles as described before^38^. Briefly, 75,000 cells were seeded at on glass coverslips and transduced with either non-targeting control or SYNJ2BP shRNA (in pLKO; Sigma Aldrich; TR30021 or TRCN0000139049, respectively) on DIV1. On DIV5 or 6, cells were washed once with PBS and fixed with 4% paraformaldehyde (PFA; Fischer Scientific; 28908)/PBS for 20 mins in room temperature (RT). Cells that were only used for immunocytochemistry were permeabilized with 0.3% Triton-X in PBS (PBST) for 10 mins in RT, blocked using 2% fetal bovine serum (Gibco; A5256801) and 2% bovine serum albumin in PBST (blocking buffer) for 60 mins in RT, and carried on with primary antibody incubation. The RNAscope protocol was adapted from the manufacturer of the probes, ACD Bio-Techne. Cells that were used for RNAscope were gradually dehydrated in 3 steps with increasing ethanol concentrations and further stored at −20°C for up to a month. Cells were rehydrated and permeabilized with 0.1% Tween 20/ PBS for 10 mins in RT. To remove endogenous peroxidase activity, the cells were then treated with hydrogen peroxide (ACD) for 10 mins in RT. After treating the cells with 1:15 diluted Protease III/PBS for 10 mins in RT, they were incubated with *Mapt* (ACD Bio-Techne; 400351-C2), RNAscope 3-plex positive control, *Ppib* (ACD Bio-Techne; 320881-C2) or RNAscope 3-plex negative control, *DapB* (ACD Bio-Techne; 320871-C1) probe for 120 mins at 40°C. Detection protocol provided by the manufacturer was followed after probe hybridization. For immunodetection following RNAscope, cells were incubated with the blocking buffer for 60 min at RT and further carried on with the primary antibody incubation.

#### Tissues

Mice were euthanized by CO_2_ over-exposure for 4 mins. They were immediately transcardially perfused with PBS. Next, the tissues were isolated and transferred to 4% PFA/PBS at 7°C overnight and then incubated in 30% sucrose/PBS at 7°C overnight. Tissues were then sectioned at 10µm thickness using a cryostat and mounted on adhesive microscopic slides (Marienfeld Histobond; 0810001). Mounted sections were either immediately stored in −80°C for later use or dried in RT for 2-4 days before performing RNAscope. Slides were baked in the oven at 60°C for 30 mins. After drawing the hydrophobic barrier slides were incubated with hydrogen peroxide for 10 mins at RT, and then with Protease III for 15 mins at 40°C. Tissues were incubated for 120 mins at 40°C with *Mapt*, or *Pink1* (ACD Bio-Techne; 536781-O1) probe. Same procedure as the cultured cells were followed for RNAscope probe detection. For immunohistochemistry after RNAscope, the slides were incubated with blocking buffer for 60 mins at RT and carried on with primary antibody incubation.

### Antibodies

All of the primary antibodies were incubated at 7°C overnight. Before secondary antibody incubation, samples were washed at least twice with PBST for 5-10 mins, each. Secondary antibody incubation was always performed at RT for a maximum of 150 mins. Both primary and secondary antibodies were diluted in blocking buffer. The following primary antibodies were used: Anti-ATP5A, mouse (1:500; Invitrogen; 43-9800), Anti-ßIII-tubulin, mouse (1:500; 2G10; Invitrogen; MA1-118), Anti-MAP2, chicken (1:1000, Novus Biologicals, NB300-213), Anti-Tau, rabbit (1:200 for ICC and 1:500 for IHC; D1M9X; Cell Signaling; 46687), Anti-RBPMS, rabbit (1:200; Proteintech; 15187-1-AP), Anti-ChAT, rabbit (1:500; EPR16590; Abcam; ab178850), Anti-SMI-31P, mouse (1:250; Biolegend; 801602). Monoclonal anti-SYNJ2a isoform specific antibodies were generated by immunization of rats with ovalbumin-coupled peptides (VFCSNSQASQPC ; aa 1238-1249; Peps4LS). Boost injections were given five and thirteen weeks later, and spleen cells were fused with myeloma cell line P3 3 63-Ag8.653 (ATCC, American Type Culture Collection) by standard procedures. Hybridoma supernatants were screened in a flow cytometry assay (iQue, Intellicyt; Sartorius) for binding to biotinylated peptides coupled to streptavidin beads (PolyAN, Berlin). Positive clones were further validated in immunoblotting and subcloned by limiting dilution to obtain stable antibody-producing monoclonal cell clones. Clone SYNJ 17H1 (rat IgG2b) was used in this study in immunostaining (1:1, hybridoma supernatant). Corresponding secondary antibodies were always diluted at 1:500.

### Microscopy and analysis of tissues

Tissues and cultured cells were mounted on Fluromount G (Invitrogen; 00-4958-02) for microscopy and were imaged in Leica STELLARIS 5 DMi8 inverted or upright confocal microscope. 63x oil 1.4 NA objective was used for immunohistochemistry and RNAscope experiments. Manually thresholded ßIII-tubulin binary masks were created in ImageJ plugin and used for extracting the axonal RNAscope or immunohistochemistry signal. “Image calculator” was used with 100*100µm squares were blindly drawn for the RNAscope channels and the density was calculated as the number of dots (each z-stack) within the defined area. For immunohistochemistry, the maximum z-projected axonal signal for Tau was normalized to that of the ßIII-tubulin. Manually thresholded binary masks were created for RBPMS and MAP2. Total area of these binary masks was calculated using “Analyze particles” and the SYNJ2a fluorescence signal was normalized to this total area (soma area in µm^2^) of RGC and MN, respectively. Two-tailed Student’s *t*-test was performed and each dot in the figures represents recordings from one tissue section.

### Microscopy and analysis of cultured cells

10x air 0.45 NA objective was used for immunocytochemistry, and 63x oil 1.4 NA objective was used for imaging RNAscope experiments, and immunohistochemistry.

#### RNAscope analysis

Mander’s correlation coefficient of individual RNAscope puncta on ATP5A-stained mitochondria was calculated using JACoP plugin within ImageJ. 10*10µm squares (avoiding nuclear region) were used to count the number of puncta per stack (in µm^2^). Maximum z-projected and digitally straightened neurites were used to calculate the Mander’s coefficient and the number of puncta per length of neurite (in µm). Each dot represents individual soma and neurite and two-tailed Student’s *t*-test was performed for these experiments.

#### Sholl analysis

100*100µm squared somato-dendritic regions, stained by MAP2, were cropped, background subtracted (50 pixels), and manually thresholded to create a mask. This mask was used to calculate the number of intersections with a step radius as 1µm (70 steps) and ramification index with the help of Sholl Analysis (Legacy Command) within ImageJ. First 60 steps are shown in the figures. Each dot in the line plot represents one biological replicate. Two-way ANOVA followed by Tukey’s HSD post-hoc tests were performed for these experiments.

#### Axon length

Simple Neurite Tracer^31^ (SNT v4.2.1) was used to trace and skeletonize primary axons (stained by SMI-31P) with longest length. Skeletonized axon lengths were calculated using the path manager graphical user interface within SNT toolbox. Two-sided Student’s *t*-test was performed for these experiments and each dot represents individual axons.

#### Tau mis-sorting

Somato-dendritic masks created using maximum z-projected MAP2 channel and axonal masks created using maximum z-projected SMI-31P channel was used to extract Tau and MAP2 fluorescence signal from somato-dendrite, and Tau and SMI-31P fluorescence signal from axons. Tau signal intensity was then separately normalized to that of MAP2 and SMI-31P. This Tau/SMI-31P ratio was further normalized to Tau/MAP2 ratio to calculate the axon/somato-dendrite distribution of Tau. Two-sided Student’s *t*-test was performed for these experiments and each dot represents individual soma and axons.

### Data analysis and statistics

Each dot shown in the quantifications involving tissues represent recordings from individual tissue sections coming from at least three animals. Each dot shown in the quantifications involving cultured neurons represent recordings from either individual cell soma or neurite/axon. Two-way ANOVA followed by Tukey’s HSD post-hoc test was performed for the Sholl analysis. Two-tailed Student’s *t*-test was performed for analyses involving 2 groups. Mean ± SEM (standard error of mean) are shown in all figures, unless otherwise mentioned. All other details are stated in the respective sub-sections.

### Other modules

ChatGPT5 was used for simplifying, creating, or looping certain R scripts. No generative AI was used for writing or modifying the text.

